# Single-cell atlas of epithelial and stromal cell heterogeneity by lobe and strain in the mouse prostate

**DOI:** 10.1101/2022.01.31.478481

**Authors:** Mindy K Graham, Roshan Chikarmane, Rulin Wang, Ajay Vaghasia, Anuj Gupta, Qizhi Zheng, Bulouere Wodu, Xin Pan, Nicole Castagna, Jianyong Liu, Jennifer Meyers, Alyza Skaist, Sarah Wheelan, Brian W Simons, Charles Bieberich, William G Nelson, Theodore L DeWeese, Angelo M De Marzo, Srinivasan Yegnasubramanian

**Affiliations:** Sidney Kimmel Comprehensive Cancer Center, Johns Hopkins University, School of Medicine, Baltimore, Maryland; Department of Neurology, Johns Hopkins University, School of Medicine, Baltimore, Maryland; Center for Comparative Medicine, Baylor College of Medicine, Houston, Texas; Department of Biological Sciences, University of Maryland at Baltimore County, Baltimore, Maryland; Department of Pathology, Johns Hopkins University, School of Medicine, Baltimore, Maryland; Department of Radiation Oncology and Molecular Radiation Sciences, Johns Hopkins University, School of Medicine, Baltimore, Maryland

**Keywords:** Single-cell genomics, Prostate, Mouse models

## Abstract

Evaluating the complex interplay of cell types in the tissue microenvironment is critical to understanding the origin and progression of diseases in the prostate and potential opportunities for intervention. Mouse models are an essential tool to investigate the molecular and cell-type-specific contributions of prostate disease at an organismal level. While there are well-documented differences in the extent, timing, and nature of disease development in various genetically engineered mouse models in different mouse strains and prostate lobes within each mouse strain, yet, the underlying molecular phenotypic differences in cell types across mouse strains and prostate lobes are incompletely understood. To address this, we examined the single-cell transcriptomes of individual mouse prostate lobes from two commonly used mouse strains, FVB/NJ and C57BL/6J. Data dimensionality reduction and clustering analysis revealed that basal and luminal cells possessed strain-specific transcriptomic differences, with luminal cells also displaying marked lobe-specific differences. Additionally, three rare populations of epithelial cells clustered independently of strain and lobe: one population of luminal cells expressing Foxi1 and components of the vacuolar ATPase proton pump (*Atp6v0d2* and *Atp6v1g3*), another population expressing Psca and other stem cell-associated genes (*Ly6a/Sca-1, Tacstd2/Trop-2*), and a neuroendocrine population expressing *Chga, Chgb*, and *Syp*. In contrast, stromal cell clusters, including fibroblasts, smooth muscle cells, endothelial cells, pericytes, and immune cell types, were conserved across strain and lobe, clustering largely by cell type and not by strain or lobe. One notable exception to this was the identification of two distinct fibroblast populations that we term subglandular fibroblasts and interstitial fibroblasts based on their strikingly distinct spatial distribution in the mouse prostate. Altogether, these data provide a practical reference of the transcriptional profiles of mouse prostate from two commonly used mouse strains and across all four prostate lobes.

## INTRODUCTION

Over the years, several genetic mouse models have been generated to study prostate cancer [1]. In contrast to the human prostate, which is divided into three glandular zones, the transition, the peripheral, and the central zones [2], the rodent prostate consists of four lobes, anterior, ventral, dorsal, and lateral. Although the dorsal and lateral lobes possess distinct differences in their histology, they are often grouped together [3,4]. Despite the anatomical and histological differences between the human and mouse prostates, animal models are an important tool for improving our understanding of prostate cancer and the changes that occur in the associated tissue microenvironment during disease development, progression, and treatment.

Prominent differences in prostate lobe and mouse strains have been recognized throughout the development of these genetic mouse models. For example, in both the Hi-MYC and Lo-MYC mouse models, the distribution of neoplastic lesions differs between the lobes, with the anterior lobe having the least [5,6]. Furthermore, there is a distinct difference in the timing of prostate cancer progression in Hi-MYC mice in an FVB background compared to those in a C57BL/6N background, which shows a significant delay in disease progression, on the order of months [7]. Likewise, in the TRAMP mouse model of prostate cancer, the C57BL/6 strain survived significantly longer than FVB mice [8]. Furthermore, tumors in the FVB TRAMP mice tended to arise in the dorsal and lateral lobes, while tumors in the C57BL/6 mice predominated in the lateral lobe [8]. Given these strain and lobe dependent differences in mouse models of prostate cancer and the known lobe-specific differences in tissue morphology and secretions [9,10], we undertook rigorous single-cell transcriptional assessments of epithelial and stromal cell types for each lobe and in two mouse strains (FVB/NJ and C57BL/6J), and assessed spatial distributions of multiple cell types using *in situ* hybridization. These analyses revealed prominent strain and lobe-specific differences in several luminal epithelial cell types, delineated the specific transcriptional programs of several epithelial, and stromal cell types that were largely similar across mouse strain and lobe, and identified novel fibroblast subclusters that displayed striking differences in their spatial distribution and lobe enrichment. Altogether, this single-cell transcriptomics study identifies the fundamental cell types that reside in the normal prostate of common mouse strains and serves as a reference to contextualize the impact of genetic alterations, exposures, or other perturbations introduced into mouse models of prostate disease.

## RESULTS & DISCUSSION

### Delineation of mouse prostate cell types reveals significant lobe and strain-specific differences in epithelial compartments

We dissected individual prostate lobes from two commonly used mouse strains, FVB/NJ and C57BL/6J (Figure 1A). Examination of the histomorphology of each lobe for each strain was consistent with previously reported lobe-specific characteristics [9] (Figure 1B). The ventral lobe of both mouse strains displayed the characteristically thin fibromuscular stroma and relatively large acini with little papillary infolding, while the lateral lobes showed distinct glandular secretions with a “chunk-like” appearance. In contrast, the dorsal lobe had relatively small acini with papillary infoldings, a thicker fibromuscular stroma, and relatively uniform secretions, consistent with prior reports. The acini of the anterior lobe had significantly more papillary infolding projections than the other lobes and were observed in both strains. Given the lobe-specific morphology of the mouse prostate, we sought to examine shared and distinct properties of each lobe using single-cell genomics.

**Figure 1.**
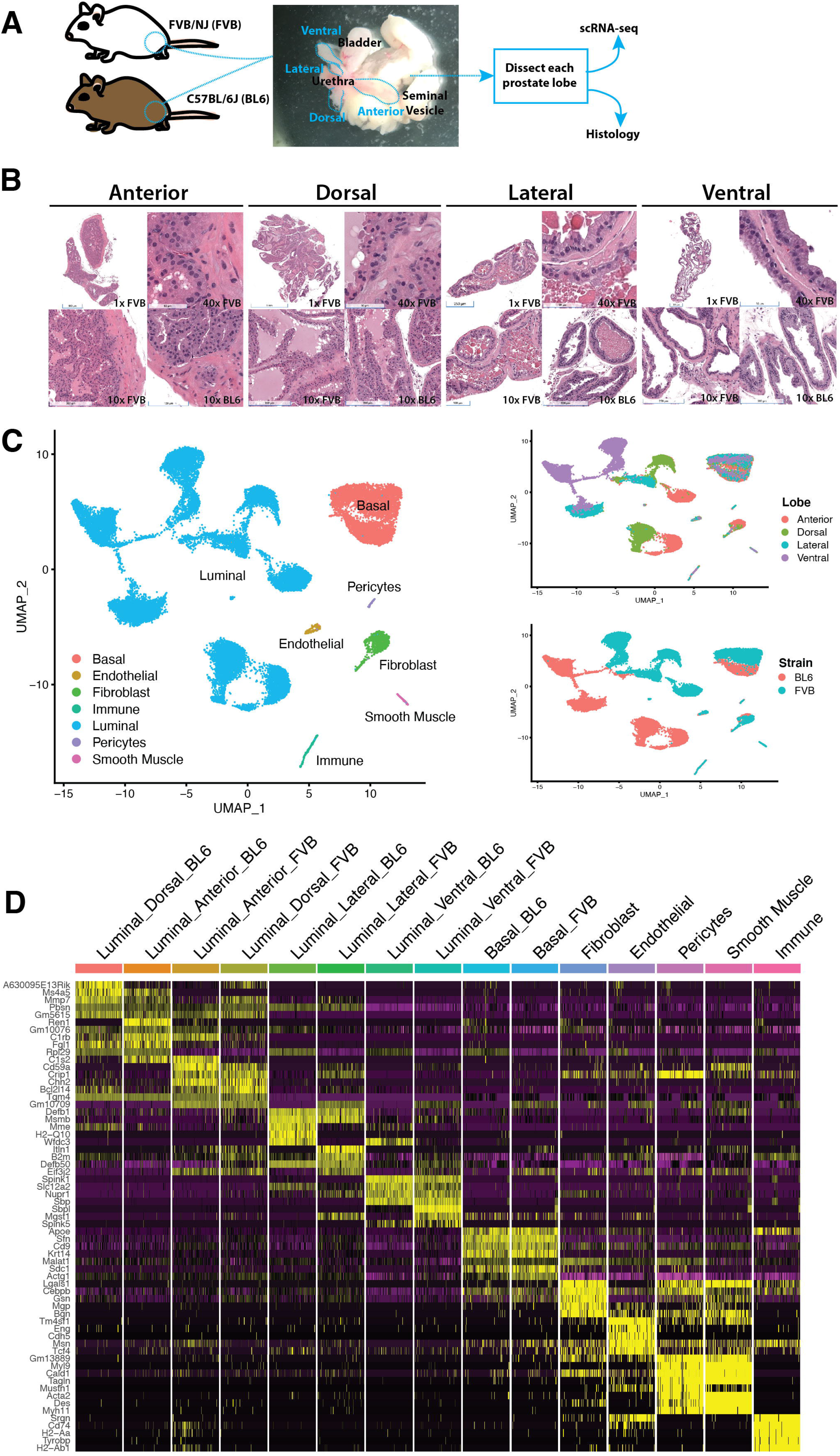
ScRNA-seq analysis of mouse prostates. A) We dissected individual prostate lobes from two commonly used mouse strains, FVB/NJ (N = 2) and C57BL/6J (N = 3), and prepared single-cell RNA-sequencing (scRNA-seq) libraries and histology for each lobe. B) Representative hematoxylin and eosin stain (H&E) staining of mouse prostate lobes from FVB/NJ (FVB) and C57BL/6J (BL6). C) Uniform Manifold Approximation and Projection (UMAP) dimensionality reduction plot of mouse prostate scRNA-seq data (27,896 cells). D) Differential gene expression across clusters visualized as a heatmap of top five genes for each cluster ranked by lowest Bonferroni adjusted p-value. Each column of the heatmap represents a cell, grouped by cluster, with 100 cells included from each group.

Single-cell RNA-sequencing (scRNA-seq) libraries were prepared separately for each strain and lobe of the mouse prostate (Table S1). The aggregated output containing 28,401 cells had 43,357 post-normalization mean reads per cell, 2,101 median genes per cell, and 8,218 median UMI counts per cell. After filtering for low quality cells, the resulting count matrix represented the expression of 18,971 genes across 27,896 cells. In general, cells clustered by cell type in uniform manifold approximation and projection (UMAP) dimensionality reduction plots (Figure 1C). Previously characterized marker genes were used to identify cell types (Figures S1, S2 & S3A), including basal and luminal epithelial, endothelial, pericyte, fibroblast, smooth muscle, and immune cell types (Figure 1C). The intermediate filament proteins *Krt8, Krt18, Krt5*, and *Krt14* were used to identify and distinguish luminal and basal epithelial cells [11,12] (Figures S1A-B and S2A-B). Stromal clusters of fibroblasts (*Pdgfra* and *Apod*)[13,14], endothelial cells (*Kdr* and *Cd93*) [15,16], and immune cells (*Cd68, Cd14 and Cd45/Ptprc*) [17,18] were identified (Figures S1C-E and S2C-E). We observed expression of smooth muscle-associated genes *Acta2* and *Tagln* [19,20] in two distinct stromal clusters (Figures S1 F&G and S2 F&G), which could be distinguished as smooth muscle cells and pericytes on the basis of expression of the pericyte-associated genes, *Rgs5* and *Notch3* [21].

The epithelial cells formed distinct subclusters, with strain- and lobe-specific luminal subclusters and strain-specific basal subclusters (Figure 1C). Differential gene expression analysis showed that multiple genes were uniquely expressed for each subcluster (Figures 1D, S3D & Table S2), suggesting the existence of strain- and lobe-specific gene expression in epithelial cell types. Prior observations of lobe-specific differences among luminal cells of the mouse prostate as assessed by single cell studies have been recently reported, but only in the C57BL/6 background [22,23]. Notably, our data revealed that multiple genes appeared to be differentially expressed between strains across epithelial cells (Figures 2A, S4 & Table S3). Some of the more prominent strain-specific genes in epithelial clusters included *Rpl129*, which was upregulated in the prostate epithelial cells of C57BL/6J mice (Figure 2A). In addition to being a component of the ribosomal 60S subunit, Rpl29 protein is also expressed on the cell surface and binds to heparin and heparan sulfate proteoglycans [24]. Epithelial cells from the prostates of FVB/NJ mice showed upregulation of the translation initiation factor, *Eif3j2*, the cyclic AMP-associated protein, *Cap1*, and the pseudogenes *Gm10260* and *Gm10709* (Figure 2A).

**Figure 2.**
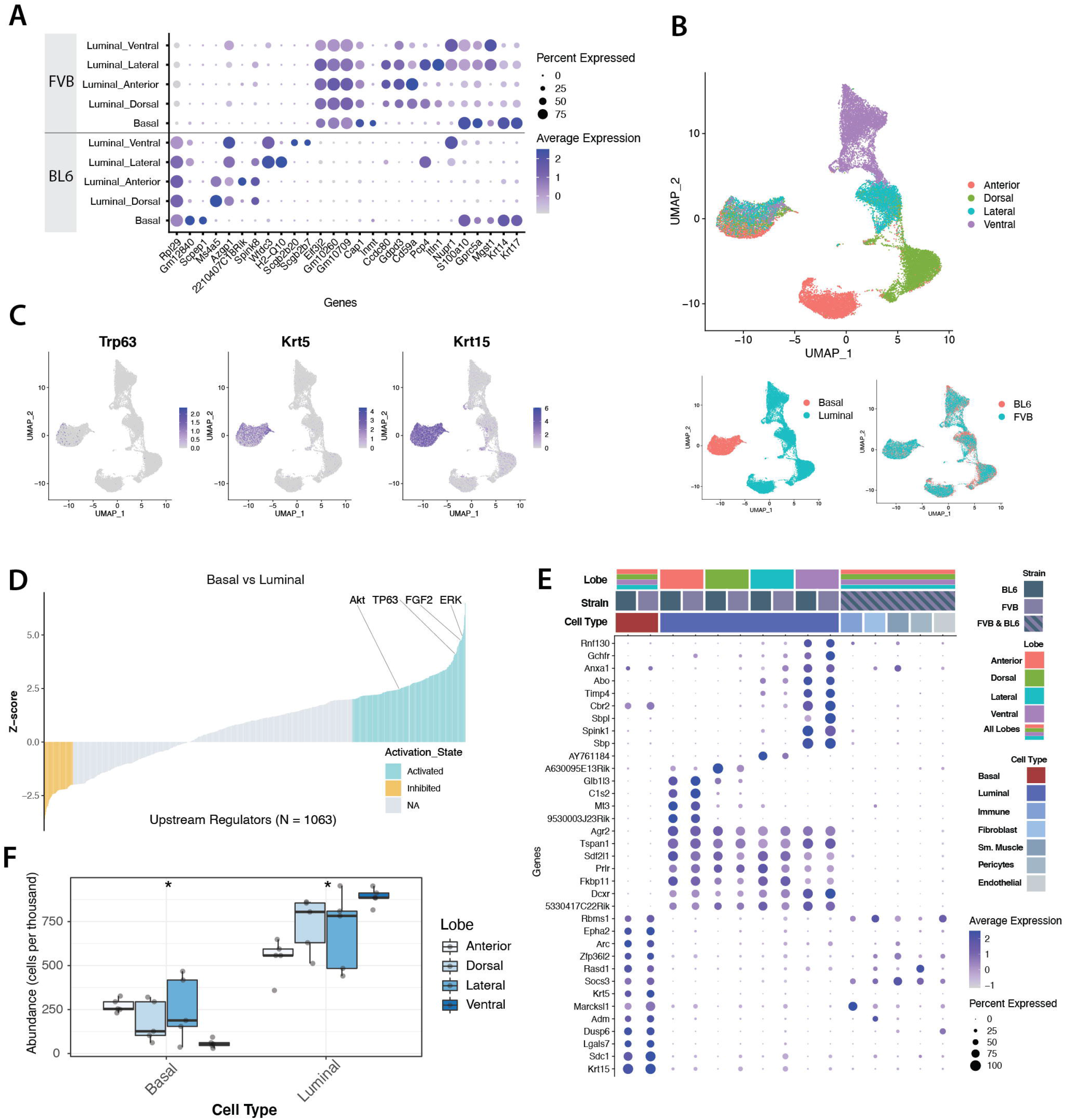
Mouse prostate epithelial cells cluster by strain and lobe. A) Dot plot of differentially expressed genes by strain in epithelial cells. For each cluster, differentially expressed genes were plotted if they had greater than two-fold change in expression, Bonferroni adjusted p-value < 0.05, and was expressed in at least 40% of cells in the cluster and less than 15% in all other clusters. B) UMAP dimensionality reduction plot of mouse prostate scRNA-seq data integrated by strain. Basal cells across all lobes partitioned together, while luminal epithelial cells clustered in four distinct groups by lobe. C) Expression of basal-associated marker genes (*Tp63*, *Krt5, Krt15)* of cells in integrated UMAP plot. The expression is assessed in Seurat (v 3.1.5) as a normalized measurement of each cell by total expression, multiplied by 10,000 and log-transformed. D) Ingenuity Pathway Analysis (IPA) of upstream regulators comparing the basal and luminal cells show that TP63, as well as basal cell maintenance associated regulators ERK, Akt and FGF2, are activated in the basal cluster. Upstream regulators with z-scores > 2 are considered activated, while z-scores < 2 are considered inhibited in basal cells. E) Dot plot of differentially expressed genes by lobe in epithelial cells. For each cluster, genes must be expressed in at least 70% of the cells and less than 20% in other cells, have a Bonferroni adjusted p-value < 0.05, and greater than two-fold change in expression. Stromal cell types are also indicated in the dot plot to show that epithelial gene expression is primarily enriched in the epithelial cell type of interest. F) Box and whisker plot of the relative abundance of basal and luminal cells for each lobe. The box represents the median and the interquartile range (IQR) with the top whisker indicating the third quartile + (1.5*IQR) and the bottom whisker indicating the first quartile-(1.5*IQR). The asterisks (*) represent an ANOVA p-value < 0.05.

We used unsupervised hierarchical clustering to capture the relative similarities or differences in the transcriptional profiles of prostate lobes from different mouse strains. The resulting dendrogram showed that the transcriptional profiles of epithelial cell types were distinct from stromal cell types, as expected (Figure S3B). Interestingly, luminal cells derived from the dorsal and anterior lobes formed a clade by strain, while the luminal cells derived from ventral and lateral lobes formed clades by lobe instead of strain. This suggests that the transcriptional profiles of dorsal and anterior lobes are the most similar, whereas the lateral and ventral lobes are each unique in ways that are shared between C57BL/6J and FVB/NJ mice. Notably, the ventral and lateral lobes shared similar histology (Figure 1B), which may in part be shaped by shared gene expression profiles. The cells originating from the ventral lobe group separated from the luminal epithelial cells originating from all other lobes in the UMAP and dendrogram plots, suggesting that ventral luminal cells are the most unique of the epithelial cell types. These analyses highlight differences and similarities between the epithelial cells in the prostates of FVB/NJ and C57BL/6J mice.

### Distinguishing features of mouse prostate lobes in luminal epithelial compartments that are conserved between strains

We performed Seurat’s integration analysis [25] in order to identify shared cell states present in both FVB/NJ and C57BL/6J strains (Figure 2B). UMAP visualization and clustering analysis of the resulting integrated data across strain revealed that basal epithelial cells across all lobes clustered together, while luminal epithelial cells clustered into four groups by lobe (Figure 2B). The basal epithelial cluster expressed the canonical markers *Krt5* and *Krt15* (Figure 2C & Table S4). While this cell cluster did also express the basal epithelial transcription factor *Tp63*, its expression was relatively low and patchy across cells in the cluster in this single cell gene expression dataset. We hypothesized that this may be due to gene dropout, which is frequently seen in single cell expression datasets [26]. To explore this hypothesis, we employed upstream regulator analysis, which can be used to infer transcription factor protein activity based on assessing the regulation of the downstream target genes of that transcription factor, even when the mRNA expression of the transcription factor itself is low or unchanging. Consistent with our hypothesis, upstream regulator analysis predicted that TP63 was one of the top activated transcription factors in the basal epithelial cluster, (Figure 2C-D & Table S5), indicating that Tp63 protein expression and signaling was likely highly active in these cells even though the degree of mRNA expression was low and patchy. Additionally, upstream regulator analysis also revealed FGF2, ERK and AKT were also activated in the basal cluster, which has been previously shown to be important in basal cell maintenance in the mouse prostate [27].

Differential expression analysis of the epithelial clusters revealed several basal and luminal-specific genes (Figure 2E). In addition to *Krt5*, a canonical prostate basal epithelial cytokeratin [11], Krt15 was also upregulated in the basal cells of both strains. The basal cell cluster also expressed genes associated with cell adhesion such as *Sdc1* [28] and *Lgals7* [29], as well as several other genes including *Dusp6, Adm, Marcksl1, Socs3, Rasd1, Zfp36l2, Arc, Epha2*, and *Rbms1*. Similarly, in luminal epithelial cells, there were numerous genes conserved across strain, as well as by lobe, including genes related to protein folding and the secretory pathway such as *Agr2* [30], *Fkbp11* [31], and *Sdf2l1* [32]; and genes encoding signal transduction proteins including *Tspan1* [33] and *Prlr* [34]. Notably, the hormone prolactin, which binds to the prolactin receptor encoded by the gene *Prlr*, is important in prostate proliferation, differentiation, and function [34].

Consistent with the lobe-specific morphology and glandular secretions conserved across mouse strains (Figure 1B), differential expression analysis showed lobe-specific gene expression as well (Figure 2E & Table S6). The lateral lobe was enriched for *AY761184*, a gene encoding a protein belonging to the family of defensin peptides, known to have antimicrobial properties [35]. The dorsal lobe expressed *A630095E13Rik (Sslp1)*, a secreted Ly-6 protein regulated by testosterone [36]. The genes conserved across strain in the anterior lobe included a component of the complement system, *C1s2;* a metallothionein protein, *Mt3*; and a hydrolase that has been reported in two gene expression signatures based on Gleason score in human prostate cancer, *Glb1l3* [37,38]. The ventral lobe, in particular, expressed a unique set of genes compared to the other lobes of the prostate, many of which encode secreted proteins, including *Anxa1*, *Abo*, *Timp4*, *Sbp*, *Sbpl*, and *Spink1* [39]. We validated lobe-specific expression of *Spink1* in the ventral lobe by *in situ* hybridization for mRNA targets (Figure 2E). Notably, the epithelial composition in the ventral lobe is also distinct compared to other prostate lobes, with far more luminal cells and fewer basal cells (Figure 2F). In the ventral lobe, luminal cells are 12-fold more abundant on average than basal cells, while the luminal cells of the other lobes range from approximately 2- to 6-fold more than basal cells.

### Rare luminal epithelial populations

Louvain graph-based clustering of epithelial cells revealed significant enrichment of individual clusters for cells originating from a particular lobe and strain. However, three rare epithelial populations clustered independently of strain and lobe (Figure 3A). One of these was a cluster of rare neuroendocrine cells (0.03%) expressing neuroendocrine-associated marker genes *Chga, Chgb*, and *Syp*. A second was a cluster of luminal cells (0.2%) expressing the forkhead transcription factor *Foxi1* and components of the vacuolar ATPase proton pump, *Atp6v0d2*, and *Atp6v1g3* (Figure 3B, Figure S5A, Table S7). Notably, Foxi1 has been reported to be necessary for the expression of multiple components of the vacuolar ATPase proton pump in epithelial cells [40]. Additionally, these *Foxi1-expressing* luminal cells also express the SERPIN serine protease inhibitors, *Serpinb6b*, and *Serpinb9.* A third cluster was comprised of progenitor-like cells (1.8%) expressing stem cell-associated genes *Ly6a/Sca-1* [41], *Tacstd2/Trop-2 [42]*, and *Psca* [43] (Figures 3B & Table S7).

**Figure 3.**
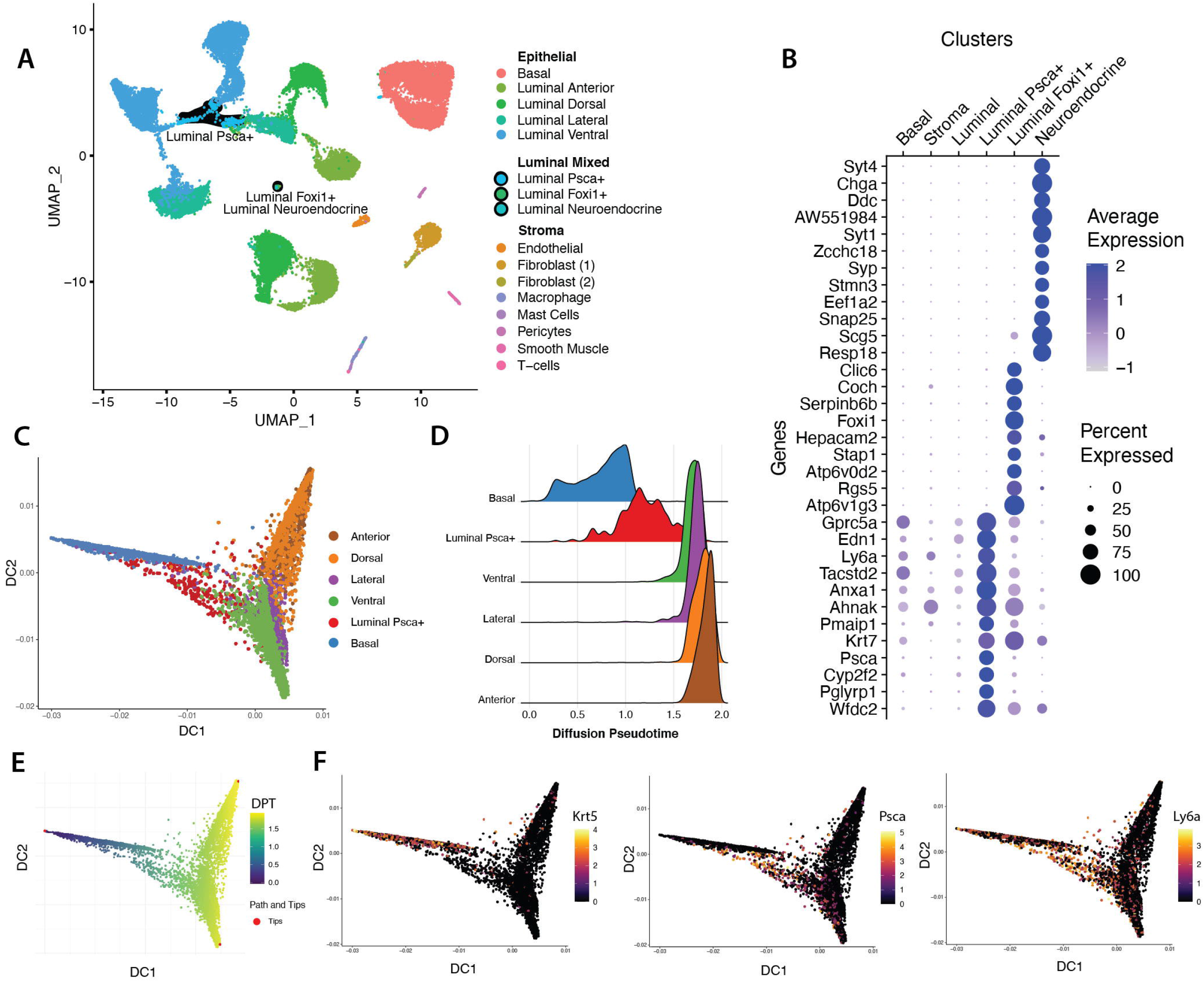
Foxi1-positive and Psca-positive luminal epithelial cells cluster independent of strain and lobe. A) Louvain clustering of cells in the UMAP plot. Highlighted are mixed populations of luminal epithelial cells that do not cluster by strain or lobe: neuroendocrine cells, *Foxi1*-positive cells, and *Psca*-positive cells. B) Dot plot of differentially expressed genes enriched in neuroendocrine cells, *Foxi1*-positive cells, and *Psca*-positive cells. For each cluster, the top 50 genes were ranked by the Bonferroni adjusted p-value and subsequently filtered for the top 15 genes by Log2 fold change. Included in the plot are basal, luminal, and stromal cells to show the specificity of gene expression in the neuroendocrine, *Foxi1*-positive, and *Psca*-positive luminal epithelial cells. C) When cells are displayed as a diffusion plot, the Psca-positive cells are positioned between basal and differentiated lobe-specific luminal epithelial cells. D) A plot of epithelial cells in diffusion pseudotime (DPT) with basal cells at time 0 and differentiated lobe-specific luminal cells at the terminal end of the scale show that Psca-positive cells are between basal and luminal cells in pseudotime. E) Diffusion plot of epithelial cells with DPT indicated. Tips indicate cells at each branch used to calculate DPT. Basal cells are at the earliest time point in DPT, progressing to anterior and dorsal as one branch and lateral and ventral as another branch. F) In the diffusion plot, Ly6a, a stem-cell-associated marker, is expressed in both Psca-positive cells and Krt5-positive basal cells.

When comparing mouse strains for the distribution of these extremely rare luminal cell populations (Table S8), we find that the neuroendocrine cells and *Foxi1* expressing cells are present in both FVB/NJ and C57BL/6J mouse strains. While there does not appear to be a strain or lobe-specific enrichment of neuroendocrine cells, Foxi1-expressing cells tend to be more enriched in FVB/NJ mice. However, with such low cell numbers represented in this dataset (< 70 cells) for these two rare cell types, it may not be possible to make interpretations for any lobe or strain-specific enrichment. Psca-expressing luminal cells were found in both strains and in every lobe, with the exception of the lateral lobe. Differential gene expression analysis with unsupervised clustering revealed that *Psca*-expressing cells were more closely related to basal epithelial cells than luminal epithelial cells, with *Psca*-positive cells expressing some basal-specific genes (Figure S5B). When diffusion mapping, a type of non-linear dimension reduction method [44], was applied to epithelial cells in pseudotime, there was a marked progression from basal to *Psca*-positive epithelial cells and then to the more differentiated lobe-specific cells, reinforcing a progenitor-like identity of these *Psca*-expressing cells (Figures 3C-E). *Krt5*, a basal-specific marker, is shown on the diffusion map and overlaps with basal cells. *Psca* is specifically expressed in the group of cells between basal and differentiated luminal epithelial cells, while *Ly6a* is expressed in both basal and *Psca*-positive luminal cells (Figure 3F). Others have also reported the existence of these rare luminal epithelial cells of the mouse prostate [22,23,45,46]. In particular, the *Psca*-expressing cells have been reported to be enriched in the proximal regions of the mouse prostate, with the lobe-specific luminal epithelial cells arising distally [22,46]. A subsequent report by Joseph et al. revealed that the *Psca*-positive cells enriched in the proximal region of the prostate extend from the urethra [47] and were determined to be the mouse counterpart of the club and hillock cells recently described in the human prostate [48].

### Stromal cell populations are primarily conserved across lobe and strain

In contrast to the epithelial cells, stromal cell clusters (smooth muscle, pericytes, and fibroblasts) and immune cells were conserved across strain and lobe, clustering by cell type and not by strain or lobe (Figure 1C, Figure S6A). Closer examination of the non-epithelial clusters revealed additional subtypes. When dimensionality reduction was restricted to stromal cell types, two distinct fibroblast populations arose, which we term subglandular and interstitial fibroblasts (see section below) (Figure 4A). Notably, in both mouse strains, one of these fibroblast populations was highly enriched in the anterior lobe in both FVB/NJ and C57BL/6J mouse strains and is orders of magnitude more abundant than any other stromal cell type (Figure 4B). As previously described, the stroma is known to display morphological differences among the lobes of the mouse prostate (Figure 1B). Potentially the enrichment of the Fibroblast 1 population may contribute to the unique appearance in the tissue morphology of the anterior lobe. In general, the mouse prostate does not appear to display strain-specific morphology, except for the fibromuscular stroma of the lateral lobe. Our data indicate that the smooth muscle cells are significantly higher in the lateral lobe of the FVB/NJ mice compared to the C57BL/6J mice (Figure 4B).

**Figure 4.**
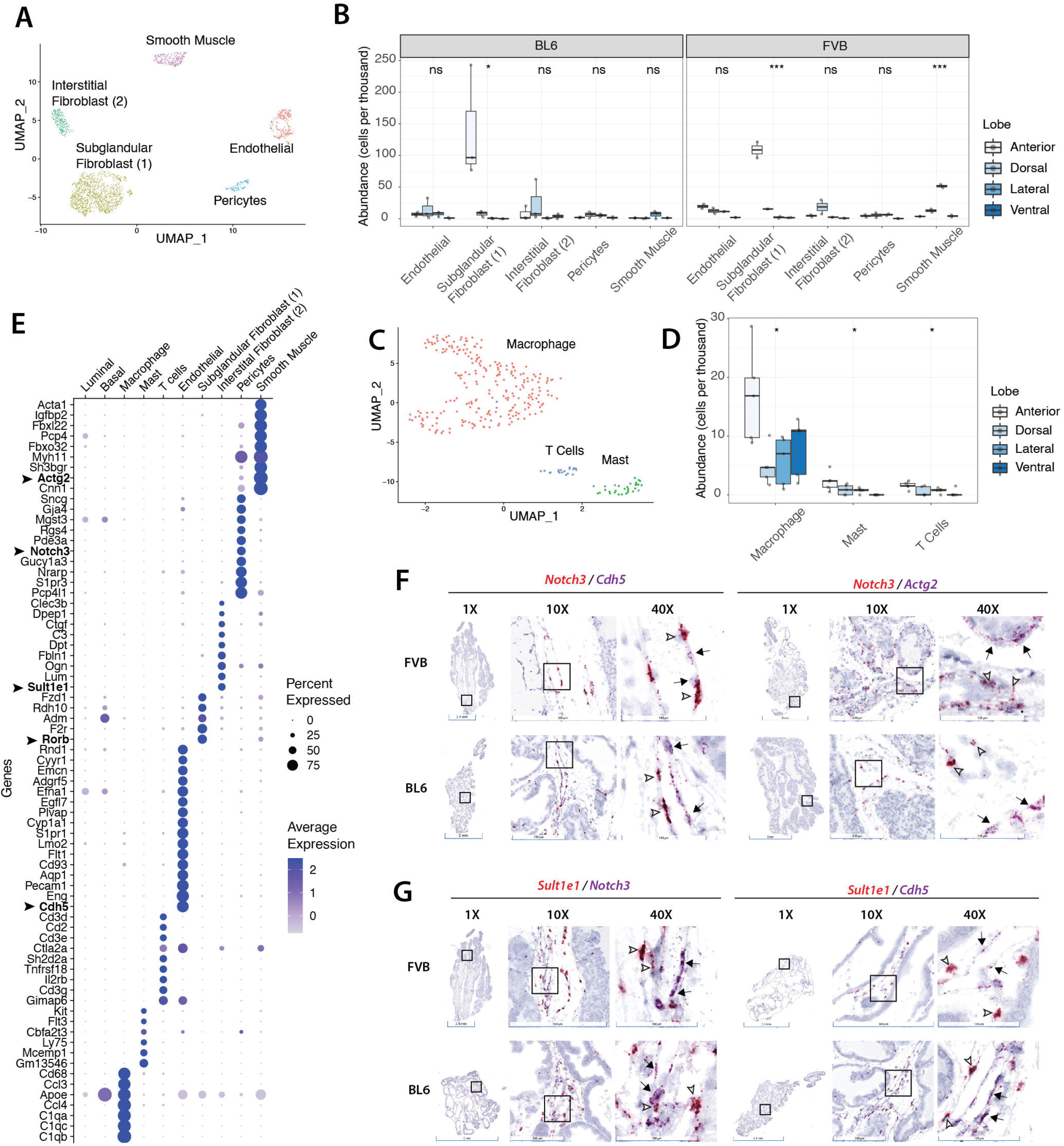
Stromal and immune cell types of the mouse prostate. A) UMAP of stromal cell types. Stromal cells include endothelial cells, smooth muscle cells, pericytes, and two distinct fibroblast populations. B) Plot of the abundance of stromal populations, including endothelial, fibroblast 1 (subglandular), fibroblast 2 (interstitial), pericytes, and smooth muscle cells in each lobe. Subglandular Fibroblast 1 cells in both C57BL/6J and FVB/NJ mice are enriched in the anterior lobe, while smooth muscle cells are enriched only in the lateral lobe of FVB/NJ mice. In the box and whisker plot, the box represents the median and the interquartile range (IQR) with the top whisker indicating the third quartile + (1.5*IQR) and the bottom whisker indicating the first quartile-(1.5*IQR). The asterisks represent an ANOVA p-value * < 0.05, and *** < 0.001, while n.s. indicates not statistically significant. C) UMAP of immune clusters. Exclusive examination of immune cells shows that cells are primarily composed of macrophages with some T-cells and mast cells. D) Plot of the abundance of immune populations, including macrophages, mast cells, and T cells in each lobe. Macrophages comprise the vast majority of immune cells in all lobes. The anterior lobe has a higher abundance of macrophages, mast cells, and T-cells than any other lobe. In the box and whisker plot, the box represents the median and the interquartile range (IQR) with the top whisker indicating the third quartile (1.5*IQR) and the bottom whisker indicating the first quartile (1.5*IQR). The asterisks represent an ANOVA p-value * < 0.05. E) Dot plot of differentially expressed genes for each stromal cluster. For each stromal cluster, the top 20 genes were ranked by the Bonferroni adjusted p-value and subsequently filtered out if >10% of other cells expressed the gene. In the case of the fibroblast 1 (subglandular) cluster, genes were filtered out if >20% of other cells expressed the gene. Luminal and basal cells are also indicated in the plot to demonstrate the specificity of gene expression. F-G) Validation of cell type-specific marker genes by multiplex (double) in situ chromogenic staining of mouse prostate lobes in C57BL/6J and FVB/NJ mice. F) *Cdh5*-expressing endothelial cells (purple chromogen, black arrow) and *Notch3-expressing* pericytes (red chromogen, open arrows, left panel) are observed at vessels, with *Actg2*-expressing smooth muscle cells (purple chromogen, black arrows, right panel) found in the fibromuscular stroma. G) *Notch3*-expressing pericytes (purple chromogen, left panel, black arrows) and *Cdh5*-expressing endothelial cells (purple chromogen, right panel, black arrows) are found juxtaposed to and in blood vessels respectively, with *Sult1e1*-expressing fibroblast cells observed in the fibromuscular stroma and near vessels (red chromogen, open arrows).

Restricting dimensionality reduction to immune cells uncovered multiple immune cell types, including macrophages, T cells, and mast cells, with macrophages as the primary constituent across all lobes (Figure 4C, Figure S6 B-C). Overall, the anterior lobe has the highest abundance of macrophages, mast cells, and T cells compared to the other lobes. In stark contrast, the ventral lobe has a lower, in some cases undetectable, amount of mast cells and T cells relative to the other lobes (Figure 4D). We assessed macrophage polarization by performing gene signature analysis using gene sets generated from two independent studies (GEO ID: GSE38705 and GEO ID: GSE161125) of stimulated and unstimulated bone marrow-derived macrophages collected from mice [49,50]. Gene signature analysis showed a progression of M0 to M1 polarization in macrophages, with a small population of M2 polarized cells. Overall, the majority of the macrophages tended towards M1 polarization (Figure S6 D). Within the T cell cluster, cells expressed *Cd8*, and markers of activated lymphocytes [51], the cytotoxins *Gzma* and *Gzmb*, and the cytokine *Ifng* (Figure S6 E). However, with limitations such as dropout in scRNA-seq, it is unclear if all cells in the T cell cluster were CD8-positive or if a portion may have been CD4-positive.

### Two fibroblast populations with distinct molecular features and localization in mouse prostate

Differential gene expression analysis revealed several cell type-specific marker genes (Figure 4E). For a subset of these genes, we validated the cell-type-specific gene expression by *in situ* hybridization for mRNA targets. *Cdh5*-expressing endothelial cells and *Notch3*-expressing pericytes were observed in vessels or juxtaposed to vessels respectively, with *Actg2*-expressing smooth muscle cells found in the fibromuscular stroma present directly adjacent to the epithelial layers (Figure 4F-G). Likewise, fibroblasts expressing the estrogen sulfotransferase *Sult1e1* were also found in the fibromuscular stroma but could also be found neighboring pericytes and endothelial cells at vessels (Figure 4G). Strikingly, the mutually exclusive *Rorb*-expressing fibroblasts and *Sult1e1*-expressing fibroblasts displayed discrete spatial patterns(Figure 5A, Figure S7). Specifically, we observed the *Rorb*-expressing Fibroblast 1 cells adjacent to the epithelial glands, similar to the Collagen IV staining pattern observed in the basement membrane of prostatic acini. In contrast, the Sult1e1-expressing fibroblasts were localized in the interstitial space between glandular acini. Based on this distinct localization, we termed the Rorb-positive Fibroblast 1 population as subglandular fibroblasts and the Sult1e1-positive Fibroblast 2 cells as interstitial fibroblasts. Recently, Joseph et al. reported observing three distinct fibroblast populations in the mouse prostate of C57BL/6J mice, which they describe as “prostate” (C3+), “urethral” (Lgr5+), and “ductal” (Wnt2+) fibroblasts [52]. The “prostate” and “ductal” fibroblasts appear to be the equivalent of our interstitial and glandular fibroblasts, respectively. Since the urethra was not intentionally included in our dissections, the “urethral” fibroblast population was likely a very rare or absent population in our dataset.

**Figure 5.**
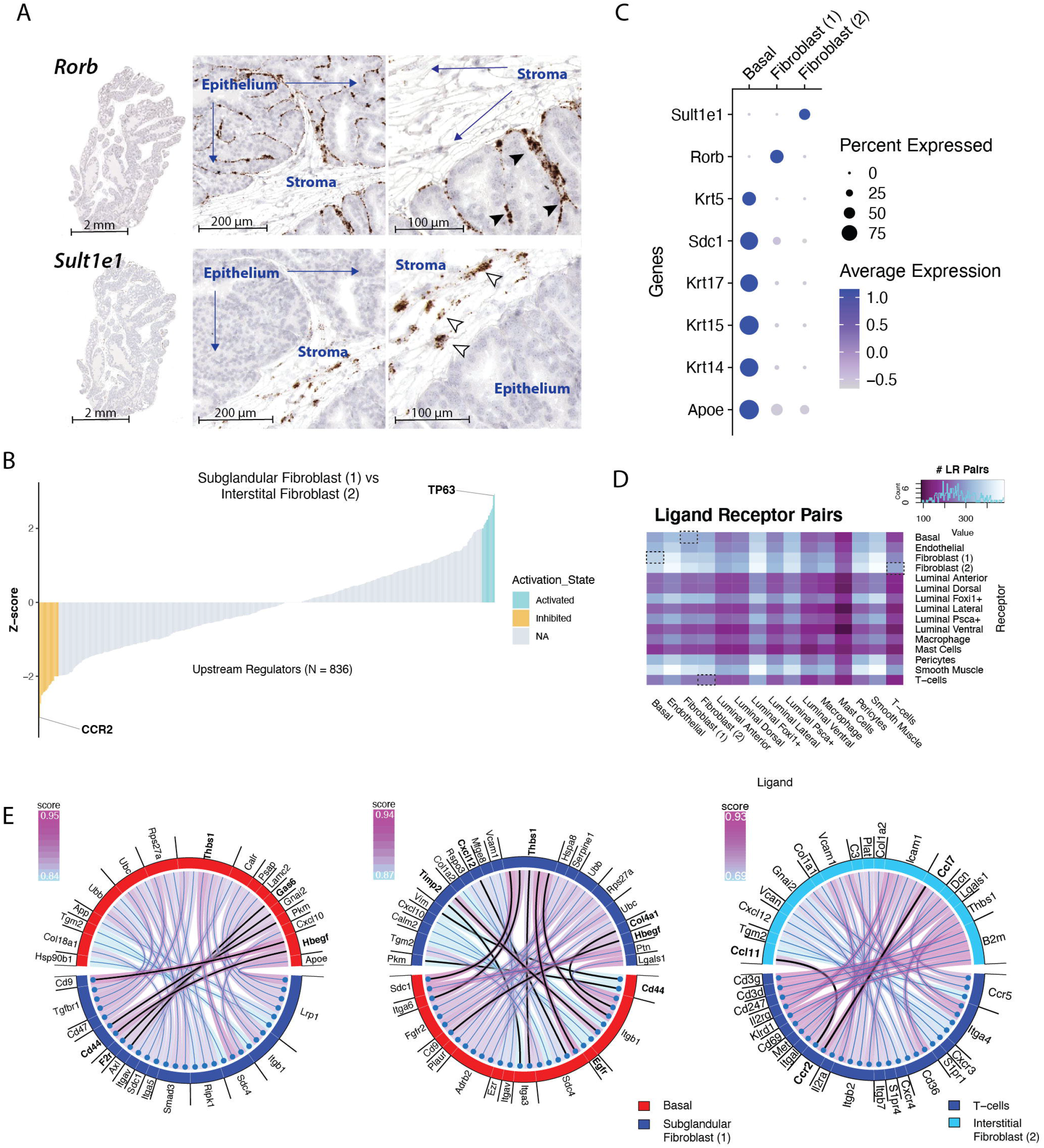
Upstream regulators of subglandular fibroblasts and interstitial fibroblasts. A) *In situ* chromogenic staining of the mouse prostate shows *Rorb*-expressing fibroblasts (black arrow heads) abutting what appears to be the basement membrane, which follows the contours of the basal aspects of the epithelial cells of prostatic acini. In contrast, *Sult1e1*-expressing fibroblasts (unfilled arrowheads) are found away from epithelial-adjacent locations in the fibromuscular stroma. This localization prompted the terms subglandular fibroblasts for the Rorb+ fibroblasts, and interstitial fibroblasts for the Sult1e1+ fibroblasts. B) IPA upstream regulator analysis comparing Subglandular (Fibroblast 1) and Interstitial (Fibroblast 2) subclusters. Upstream regulators with z-scores > 2 are considered activated while z-scores < 2 are considered inhibited in subglandular fibroblast cells. Note that the inhibited upstream regulators are activated in interstitial Fibroblast 2 cells. Highlighted is Ccr2, which is upregulated in the interstitial fibroblast 2 cluster compared to the subglandular fibroblast 1 cluster. C) Dot plot shows that *Rorb*-expressing subglandular Fibroblast 1 cells and *Sult1e1*-expressing interstitial Fibroblast 2 cells do not express Basal-specific marker genes. D) Matrix heatmap showing the number of ligand-receptor pair interactions for each cluster. Dotted boxes indicate subglandular Fibroblast 1 with Basal cell interactions, and interstitial Fibroblast 2 with T cell interactions. E) Chord plots of the top ligand and receptor interactions between basal cells and subglandular Fibroblast 1 cells (first and middle panel), with downstream targets of Tp53 bolded. The right chord plot shows ligand (interstitial Fibroblast 2) and receptor (T cells) interactions, with Ccr2 associated ligands in bold.

In addition to expressing genes associated with retinoic acid signaling, *Rorb*, and *Rdh10*, the subglandular fibroblasts expressed the Wnt receptor, *Fzd1* (Figure 4E). Upstream regulator analysis revealed fundamentally distinct pathways, with subglandular fibroblasts having activation of the DNA binding protein Tp63 (Figure 5B). Tp63 is an epithelial-specific gene, and in the prostate, it is exclusively expressed in basal epithelial cells [53]. While the subglandular fibroblast cells did not express basal cell-specific genes (Figure 5C), ligand-receptor pair analysis inferred more than 200 potential interactions between basal and subglandular fibroblast cells (Figure 5D). A subset of those inferred interactions included downstream targets of Tp63, including the cell surface glycoprotein Cd44, the thrombin receptor F2r, the metallopeptidase inhibitor Timp2, and a major structural component of the basement membrane, Collagen Type IV Alpha 1 Chain Col4a1 (Figure 5E, first and second panels). Notably, subglandular fibroblast cells neighbor the perimeter of the epithelial glands where basal cells reside in the prostate (Figure 5A, Figure S7), suggesting that the close proximity of these cell types and the numerous potential ligand-receptor interactions may account for the upregulation of Tp63 activity observed in the subglandular fibroblast cells.

Despite sharing many genes with the subglandular fibroblast cells (Figure S5A), interstitial fibroblasts showed upregulation for genes encoding extracellular matrix proteins, *Lum*, *Ogn, Fbln1*, and *Dpt* (Figure 4E). Pathway analysis comparing both fibroblast clusters implicated the fibrosis-associated chemokine receptor, *Ccr2* [54], as one of the top upstream regulators in interstitial fibroblasts (Figure 5B). Interestingly, examining ligand-receptor pairs by expression among all cell clusters indicated that T-cell ligands paired the most with smooth muscle and interstitial fibroblast receptors (Figure 5D). In particular, one of the highest ligand-receptor scores between T cells and interstitial fibroblasts included the chemokine receptor Ccr2 and the cytokines Ccl7 and Ccl11, suggesting that interstitial fibroblasts cells may directly engage with infiltrating or tissue resident immune cells in the tissue microenvironment.

## Conclusions

In these foundational single-cell studies of strain and lobe-specific differences in the mouse prostate, we have uncovered previously uncharacterized cell types and nominated unique molecular markers of multiple cell types for a more granular *in situ* examination of mouse prostate tissues. Our scRNA-seq analysis of normal mouse prostates revealed that lobe and strain-specific differences are primarily restricted to epithelial cell types. Given the morphological differences of mouse prostate lobes (Figure 1B), it is perhaps not entirely unexpected that luminal epithelial cells would cluster separately by lobe in our analysis (Figure 1C). However, the strain-specific clustering of both basal and luminal epithelial cells is noteworthy, and to the best of our knowledge, has not been previously reported elsewhere. This striking observation suggests that transcriptional differences among the basal and luminal epithelial cells from different mouse strains are pronounced enough to drive strain-specific clustering. Future studies interrogating genomic and transcriptomic sequencing data for expression quantitative trait loci (eQTL) analysis [55] may be informative in revealing the primary contributors of these strain-specific differences. Notably, the molecular consequences of genetic variation between mouse strains is cell type-specific in the prostate, as only the basal and luminal cells clustered by strain in our analysis. Given these strain-specific differences observed in the epithelial cells of the mouse prostate, such inter-individual differences also potentially exist in human prostate luminal and basal cells and may contribute to disease risk. Such inter-individual variation may therefore modify the phenotypic effects of driver mutations and contribute to the observed heterogeneity in disease aggressiveness and therapeutic response/outcomes.

Although the majority of luminal cells clustered by strain and lobe in mouse prostates, we also observed three rare luminal populations that clustered independently of lobe and strain: neuroendocrine cells, *Foxi1* expressing cells, and *Psca* expressing cells. Pseudotime analysis of epithelial cells showed a clear progression from basal to *Psca*-positive epithelial cells and then to the more differentiated lobe-specific cells, suggesting a progenitor-like identity of these *Psca*-expressing cells.

Stromal cell types were largely conserved across strain and lobe. We found that the ventral lobe had a distinctly higher luminal-to-basal cell ratio and fewer mast cells and T cells than other lobes of the mouse prostate (Figures 2F & 4D). In contrast, the anterior lobe was uniquely enriched for the subglandular fibroblast population, which comprised the vast majority of the stroma and had more immune cells relative to other lobes (Figures 4B & D). On the other hand, interstitial fibroblasts tended towards immune-associated interactions, with Ccr2 being the most prominent (Figures 4B & E). Perhaps the most striking observation was the discrete spatial patterns in the mouse prostate of these mutually exclusive *Rorb*-expressing subglandular fibroblasts *Sult1e1*-expressing interstitial fibroblasts and the distinct expression programs of each subtype. Overall, these findings suggest an intriguing possibility that these fibroblast subtypes may have different biological roles in the tissue, with subglandular fibroblasts engaging the epithelial cells of prostatic acini and interstitial fibroblasts mediating immune-associated interactions.

Taken together, the combination of lobe-specific differences in luminal cells and the stromal composition in the prostate may contribute to the histological differences observed between mouse prostate lobes. Furthermore, the strain and lobe-specific differences in epithelial cells and immune cells may, in part, drive the intrinsic propensity for the ventral, dorsal, and lateral lobes to develop prostatic lesions (e.g. prostatic intraepithelial neoplasia and adenocarcinomas) more rapidly than the anterior lobe in the MYC, TRAMP, and PTEN mouse models of prostate cancer [5-8,56]. Immune infiltrating cells, particularly myeloid-derived suppressor cells at pre-cancerous lesions, have been implicated in the strain-specific differences in the Hi-MYC prostate cancer model [7]. Our data indicate that, at least in the wild-type setting, there are no significant differences in the composition of the immune cell types between the FVB/NJ and C57BL/6J mouse strains. The myeloid cell types we detected in mouse prostates included mast cells and macrophages with a tendency towards M1 polarization. Overall, the findings of this study help establish the fundamental cell types residing in the normal mouse prostate of common mouse strains and serve as a reference to better understand how pharmacological or genetic mouse models of prostatic disease can be influenced by the normal biology of cells in the prostate.

## MATERIALS AND METHODS

### Mouse models and prostate dissection

Wildtype C57BL/6J (N = 3) and FVB/NJ (N = 2) mice were purchased from Jackson Laboratory and were maintained until they reached six months of age. Individual prostate lobes were dissected, and single-cell RNA-sequencing (scRNA-seq) libraries for each lobe were prepared separately. Mice were euthanized using carbon dioxide asphyxiation, and the urogenital tract was removed and placed into a petri dish containing 50 ml of Hanks’ Balanced Salt Solution (HBSS, Gibco 14175-079). Under a dissection microscope, adipose tissues were removed from the urogenital tract to isolate the prostate. The four pairs of lobes (anterior, dorsal, lateral, and ventral lobes) were separated and dissected from the urethra by forceps. Half of each lobe was used for single-cell RNA-Seq, a quarter was formalin-fixed & paraffin-embedded, and the remaining quarter was frozen.

### Dissociation of mouse prostate

Mouse prostate tissues were minced with razor blades and then digested in 0.25% Trypsin-EDTA (Gibco 25200-072) for 10 min at 37°C, followed by incubation for 2.5 hours at 37°C with gentle agitation in DMEM containing 10% FBS, 1 mg/mL Collagenase Type I (Gibco 17100-017), and 0.1 mg/mL of DNase I (Roche 10104159001). Digested tissues were centrifuged at 400 x g for 5 minutes and subsequently washed with HBSS and further incubated in 0.25% Trypsin-EDTA for 10 min at 37°C. Cells were suspended in DMEM containing 10% FBS and 0.4 mg/mL of DNase I. Cell aggregates were dissociated by pipetting repeatedly. Cells were then passed through a 40 μm cell strainer to generate a single-cell suspension. An aliquot of cells was stained with trypan blue (Gibco 15250-061) and counted using a hemocytometer to assess cell yield and viability.

### Single-cell RNA-sequencing and data pre-processing

Libraries for scRNA-seq were prepared using the 10x Genomics Chromium Single Cell 3’ Library and Gel bead Kit V2 (CG00052_RevF) according to the manufacturer’s protocol for each dissected prostate lobe. The cDNA libraries were subsequently sequenced (150 bp paired-end) on the Illumina HiSeqX platform. Each sequenced library was demultiplexed to FASTQ files using Cell Ranger (10x Genomics). Cell Ranger (version 2.2.0) count pipeline was used to align reads to the mm10 mouse transcriptome and create a gene by cell count matrix. Cell Ranger (version 2.2.0) aggr pipeline was used to aggregate the counts across all the cDNA libraries generated for each mouse prostate lobe. Sequencing files can be accessed on NCBI GEO (GSE165741). Seurat (version 3.1.5) [25] was used to further pre-process the data. Cells were discarded if they contained counts of less than 200 or greater than 4,500 genes to minimize contamination of RNA-containing supernatants or multiplets. Cells with mitochondrial gene percentages exceeding 10% were also discarded.

### Dimensionality reduction & cell type identification

Seurat (version 3.1.5) was used to normalize, scale, and log-transform the count data [25,57]. Principal components were computed based on the 2,000 most variable genes in the data set. The first 20 principal components were used to compute the Uniform Manifold Approximation & Projection (UMAP) dimensions and perform Louvain clustering at a resolution of 0.4. Cluster-specific marker genes were identified for each of the resulting clusters using differential expression analysis. Clusters were assigned cell type identities using previously characterized marker genes, specifically *Pdgfra* and *Apod* for fibroblasts, *Kdr* and *Cd93* for endothelial cells, *Cd68, Cd14, Cd45/ Ptprc* for immune cells, and *Rgs5* and *Notch3* for pericytes. Smooth muscle cells were identified by the presence of *Acta2* and *Tagln* and the absence of pericyte markers. Basal cells were identified using the markers *Krt5* and *Krt14*, while luminal clusters were identified based on the presence of *Krt8* and *Krt18* and the absence of *Krt5* and *Krt14*. Luminal cells that clustered independent of strain and lobe were identified as *Psca*-expressing luminal and *Foxi1*-expressing luminal cells. Embedded within the cluster of *Foxi1*-expressing luminal cells were rare *Chga/Syp*-expressing cells. Any cells within this cluster expressing *Chga/Syp* were annotated as neuroendocrine cells in the seurat object. In a subsequent cluster analysis, the data were integrated to minimize variability between FVB/NJ and C57BL/6J while observing differences in cell types. In the integration step, 2000 anchor features were chosen and canonical correlation analysis was selected for dimensionality reduction with 20 dimensions used to specify the neighbor search space, and 200 neighbors (k-filter) were specified for anchor filtering. When dimensionality reduction was restricted to stromal cell types, excluding immune cells, 20 principal components were used to compute the UMAP dimensions with Louvain clustering at a resolution of 0.1. Similar parameters were selected for cluster analysis restricted to immune cells, with Louvain clustering performed at a resolution of 0.3.

### Differential gene expression analysis

Seurat (version 3.1.5) was used to perform differential gene expression analyses of all clusters. Genes considered in the analysis needed to be expressed in at least 25% of cells in a cluster with a minimum of natural log fold change of 0.25. Outputs of differentially expressed genes are included as supplemental tables, limited to genes with Bonferroni corrected p-value < 0.05 and fold change expression > 2. The gene filtering parameters used to generate each heatmap and dot plot from differential gene expression analysis are described in the legend of each figure.

### Upstream Pathway analysis

Upstream regulators that are likely to mediate the observed gene expression differences across cell clusters were analyzed using Ingenuity Pathway Analysis (IPA) [58]. Biological networks constructed from known interactions in the published literature were used to infer upstream molecular regulators. Using observed differential gene expression, a z-score was derived from predicted up or down-regulation of relevant genes in a pathway. Seurat was used to perform differential gene expression analyses comparing basal to luminal clusters, and subglandular (Fibroblast 1) to interstitial (Fibroblast 2) clusters, testing genes with a minimum of 10% of cells detected in either population and a natural log fold change > 0.1. The subsequent IPA analysis was implemented with differentially expressed genes with at least a two-fold change and a false discovery rate < 0.05. The predicted transcription factors in the epithelial comparison and the predicted upstream regulators in the fibroblast comparison were plotted with their associated z-scores.

### Trajectory analysis

The counts matrix for previously defined basal and luminal epithelial cell clusters were manually scaled to reads per million (RPM) and log10-transformed. Diffusion components and diffusion pseudotime (DPT) parameters were computed using Destiny (version 3.1.1) [44].

### Multiplex Chromogenic in situ hybridization (CISH) staining

A subset of cell type specific marker genes were validated by CISH using the RNAscope 2.5 HD Reagent kit-Brown (Cat.No 322300, ACD) and RNAscope 2.5 HD Duplex Reagent Kit (Cat. no 322430, ACD). Probes included *Rorb* (Mm-Rorb C1, Cat# 44427-C1), *Sult1e1* (Mm-Sult1e1 C1 and C2, Cat# 900181), *Actg2* (Mm-Actg2 C1, Cat# 483811-C1), *Notch3* (Mm-Notch3 C1 and C2, Cat# 425171) and *Cdh5* (Mm-Cdh5 C1, Cat# 312531-C1). To prepare slides for CISH staining, FFPE slides were baked for 30 minutes at 60°C, and deparaffinize by incubating slides at room temperature (RM) for 10 minutes in xylene twice and then subsequently incubating in 100% ethanol twice, and finally left to air dry. Hydrogen peroxide solution was added onto the slides for 10 minutes at RT. Slides were steamed in 1 x RNAscope Target retrieval reagent at 100°C for 18 minutes, followed by protease plus digestion for 30 minutes at 40°C to allow target accessibility. The C1 probe was used for single CISH staining, while both C1 and C2 probes were used for duplex CISH staining. Probes were mixed at a 1:50 ratio, added to slides, and incubated in the HybEZ TM Oven for 2 hours at 40°C. Signal amplification and detection were performed according to the manufacturer’s protocol. The C1 probe signal was detected with DAB or Discovery Purple (Cat No 7053983001, Ventana), and the C2 signal was detected with ImmPACT Vector red (SK-5105). Slides were counter-stained with 50% Gill’s Hematoxylin for 2 min, rinsed in 0.02% ammonia water for 10 seconds, treated in 100% ethanol and xylene and cover slipped with Cytoseal mounting medium.

### Ligand-receptor pair analysis

Data preprocessed and integrated by mouse strain in Seurat (version 3.1.5) were extracted and analyzed in SignalCellSignalR (version 1.2.0) [59] to infer ligand-receptor (LR) pair interactions of each cluster based on the normalized expression. Paracrine interactions and autocrine interactions were independently assessed and the outputs were subsequently combined with any overlapping duplex LR pairs filtered such that only unique interactions remained. Interaction pairs with LR scores > 0.5 were considered in the generation of the matrix heatmap. Chord diagrams were generated from the top 30 interactions between ligands and receptors of Fibroblast 1 and Basal cells, and LR pairs of Fibroblast 2 and T cells.

### Gene signature analysis of immune cell clusters

The tendency for M1 or M2 polarization of macrophages was examined using AUCell (version 1.12.0) [60]. An expression matrix containing the cell by gene counts was used to build gene expression rankings for each cell. The area under curve (AUC) enrichment was assessed from gene sets generated from publicly available datasets. The threshold used to calculate the AUC was based on the top 5% of genes in the gene expression rankings for each cell. As a control, a random selection of 500 genes was used to generate the Random gene set. The GSE38705 M0 and GSE38705 M1 gene sets were generated from the Hybrid Mouse Diversity Panel of macrophages (GEO ID: GSE38705)[49]. GEOR2 was used to compare the expression of all baseline macrophage samples collected from 92 inbred mouse strains (M0, N = 172) to bacterial lipopolysaccharide treated macrophages (M1, N = 182). The GSE38705 M0 gene set contains all genes with Log2 FC > 1 and Benjamini & Hochberg FDR-adjusted p-value < 0.05. The GSE38705 M1 gene set contains all genes with Log2 FC < −1 and Benjamini & Hochberg FDR-adjusted p-value < 0.05. The GSE161125 M0, GSE161125 M1, and GSE161125 M2 gene sets were derived from a scRNA-seq dataset of bone marrow-derived macrophages collected from wild-type C57BL/6J mice (GEO ID: GSE161125) [50]. Macrophages in culture media (M0, N = 1914 cells), culture media containing bacterial lipopolysaccharide and interferon-gamma (M1, N = 1815 cells), or culture media containing in interleukin 4 (M2, N = 1474 cells) were analyzed in Seurat (version 3.1.5). Marker genes for each group (M0, M1, and M2) were determined based on the following criteria, at least 25% of cells expressed the gene, and the gene had a natural log fold change > 0.25 as determined by differential gene expression analysis. The marker genes for M0, M1, and M2 were used to generate the GSE161125 M0, GSE161125 M1, and GSE161125 M2 gene sets. All gene sets used for the gene signature analysis are listed in Table S8. For each gene set, the top 150 cells in the entire dataset (N = 28,399 cells) with the highest AUC scores were marked as positively enriched, although the associated UMAP plots only display the immune cell clusters in order to highlight macrophage enrichment.

## Supporting information

Sigure S6

## ACKNOWLEDGEMENTS

We thank the members of the Sidney Kimmel Comprehensive Cancer Center’s Experimental and Computational Genomics Core, supported by Cancer Center Support Grant P30CA006973, for support with the single cell sequencing studies and data pre-processing and analysis. This work was supported by NIH/NCI grants P50CA058236, U01CA196390, P01CA247886, and by the Prostate Cancer Foundation, The Allegheny Health Network Johns Hopkins Pilot Project Grant, The Patrick C. Walsh Fund, The Irving Hansen Foundation, and the Maryland Cigarette Restitution Fund.

