## Supplemental Materials for "Single-cell atlas of epithelial and stromal cell heterogeneity by lobe and strain in the mouse prostate"

SUPPLEMENTARY MATERIALS

SUPPLEMENTARY TABLES

**Table S1. Cell totals from mouse prostates by strain, lobe, and cell type.**

**Table S2. List of genes differentially expressed for each cluster by cell type, strain, and lobe.**

Pct. 1 indicates % cells the gene is detected in the cluster. Pct.2 indicates % cells the gene is detected in all other clusters. The adjusted p-value is based on Bonferroni correction. At least 25% of cells must express the gene and have a minimum log fold change of 0.25. List contains genes with Bonferroni corrected p-value < 0.05 and fold change expression > 2.

**Table S3. List of genes differentially expressed by strain in epithelial cells.**

Pct. 1 indicates % cells the gene is detected in the cluster. Pct.2 indicates % cells the gene is detected in all other clusters. The adjusted p-value is based on Bonferroni correction. At least 25% of cells must express the gene and have a minimum log fold change of 0.25. List contains genes with Bonferroni corrected p-value < 0.05 and |log2FC| > 1. Positive log2FC indicates positive differential expression in BL6, while negative log2FC indicates positive differential expression in FVB.

**Table S4. Differentially expressed genes in basal vs luminal cell types.**

Pct. 1 indicates % cells the gene is detected in the cluster. Pct.2 indicates % cells the gene is detected in all other clusters. The adjusted p-value is based on bonferroni correction. At least 25% of cells must express the gene and have a minimum of log fold change 0.25. List contains genes with Bonferonni corrected p-value < 0.05 and |log2FC| > 1. Positive log2FC indicates positive differential expression in basal cells while negative log2FC indicates positive differential expression in luminal cells.

**Table S5. Ingenuity Pathway Analysis (IPA) of upstream regulators in basal versus luminal cell types.**

List of all transcription regulators with calculated Z-scores. Positive Z-scores indicate activation in basal cells, while negative Z-scores indicate activation in luminal cells.

**Table S6. Lobe-specific differentially expressed genes independent of strain.**

Pct. 1 indicates % cells the gene is detected in the cluster. Pct.2 indicates % cells the gene is detected in all other clusters. The adjusted p-value is based on bonferroni correction. At least 25% of cells must express the gene and have a minimum of log fold change 0.25. List contains genes with Bonferroni corrected p-value < 0.05 and |log2FC| > 1.

**Table S7. List of genes differentially expressed for each cluster by Louvain clustering.** Pct. 1 indicates % cells the gene is detected in the cluster. Pct.2 indicates % cells the gene is detected in all other clusters. The adjusted p-value is based on bonferroni correction. At least 25% of cells must express the gene and have a minimum of log fold change 0.25.

**Table S8. Cell totals of rare luminal cell types from mouse prostates by strain and lobe.** Percent is calculated by dividing cell number by total cell number of all the mouse lobes.

**Table S9. List of genes used for AUC geneset analysis**

SUPPLEMENTARY FIGURES AND LEGENDS

**Figure S1. UMAP plots of canonical marker gene expression used to identify cell types of clusters.**

The expression is assessed in Seurat (v 3.1.5) as a normalized measurement of each cell by total expression, multiplied by 10,000 and log-transformed. Cell types identified include A) luminal expressing *Krt8* and *Krt18*, B) basal expressing *Krt5* and *Krt14*, C) fibroblast expressing *Pdgfra* and *Apod* , D) endothelial expressing *Kdr* and *Cd93*, E) immune expressing *Cd68* and *Cd14*, F) pericytes expressing *Rgs5* and *Notch3*, and G) smooth muscle expressing *Acta2* and *Tagln*.

**Figure S2. Violin plots of canonical marker gene expression in all clusters that group by cell type.**

Clusters that are grouped by cell type include luminal, basal, fibroblast, endothelial, immune, pericytes and smooth muscles. Genes include A) *Krt8* and *Krt18*, B) *Krt5* and *Krt 14*, C) *Pdgfra* and *Apod*, D) *Kdr* and *Cd93*, E) *Cd68* and *Cd14*, F) *Rgs5* and *Notch3*, and G) *Acta2* and *Tagln*.

**Figure S3. Marker gene expression of identified cell types.**

A) Dotplot of marker genes of all identified cell types. B) Heatmap of differentially expressed genes for each UMAP cluster by cell type, strain, and lobe. Genes have a Bonferroni adjusted p-value < 0.05 and gene expression fold change > 2. Refer to Table S2 for a complete list of genes.

**Figure S4. Strain specific genes of mouse prostate.**
Venn diagrams of strain specific genes for basal cells and luminal cells by lobe in A) C57BL/6J and B) FVB/NJ. Refer to Table S3 for a complete list of genes.

**Figure S5. Mixed luminal epithelial populations cluster independent of strain or prostate lobe.** A) Differential gene expression across Louvain clusters visualized as a heatmap of top five genes for each cluster ranked by lowest Bonferroni adjusted p-value. B) Heatmap of differentially expressed genes for each group determined by Louvain clustering with unsupervised hierarchical clustering. Genes have a Bonferroni adjusted p-value < 0.05 and gene expression fold change > 2. Refer to Table S7 for a complete list of genes.

**Figure S6. Characterization of immune cell clusters.** A) UMAP of immune clusters marked by strain (middle panel) and lobe (right panel) show a mixed distribution. Immune cell types identified by marker gene expression B) of UMAP clusters, including macrophages (*Cd68*, *Cd14*), mast cells (*Kit*, *Mcemp1*), and T cells (*Cd3g*, *Cd2*). C) Violin plots of marker gene expression of immune clusters. D) Macrophage polarization gene set analysis (AUCell v 1.12.0). AUCell histograms show area under curve (AUC) scores of all cells for each gene set (Random, GSE38705 M0, GSE38705 M1, GSE161125 M0, GSE161125 M1, GSE161125 M2). AUC cell cut offs of top 150 cells for each gene set for all cells. UMAP of immune cell clusters highlight cells greater than the AUC score cut off in blue, and gene set activity indicated as red (high gene set enrichment) and light pink (low gene set enrichment). E) Heatmap of select T cell associated genes for all cell clusters. Each column represents each cell type downsampled to 25 cells.

**Figure S7. Subglandular and interstitial fibroblasts**
Multiplex in situ chromogenic staining of the anterior lobe (AP) shows *Rorb*-expressing subglandular fibroblasts (brown chromogen, black arrowheads). These cells are localized in a manner in which they appear to be closely abutting the basement membrane, which follows the contours of the basal aspects of the epithelial cells of prostatic acini. By contrast, the *Sult1e1*-expressing interstitial fibroblasts (red chromogen, open arrowheads) are found away from epithelial-adjacent locations in the fibromuscular stroma. The ventral lobe (VP) has little to no

*Rorb*-expressing subglandular fibroblasts (bottom second and fourth panels). The right panel shows Collagen IV IHC staining of the FVB/NJ mouse prostate lobe, which highlights the basement membrane (black arrows).
