## Supplementary figures and images for "Single-cell atlas of epithelial and stromal cell heterogeneity by lobe and strain in the mouse prostate"

### Figure S1

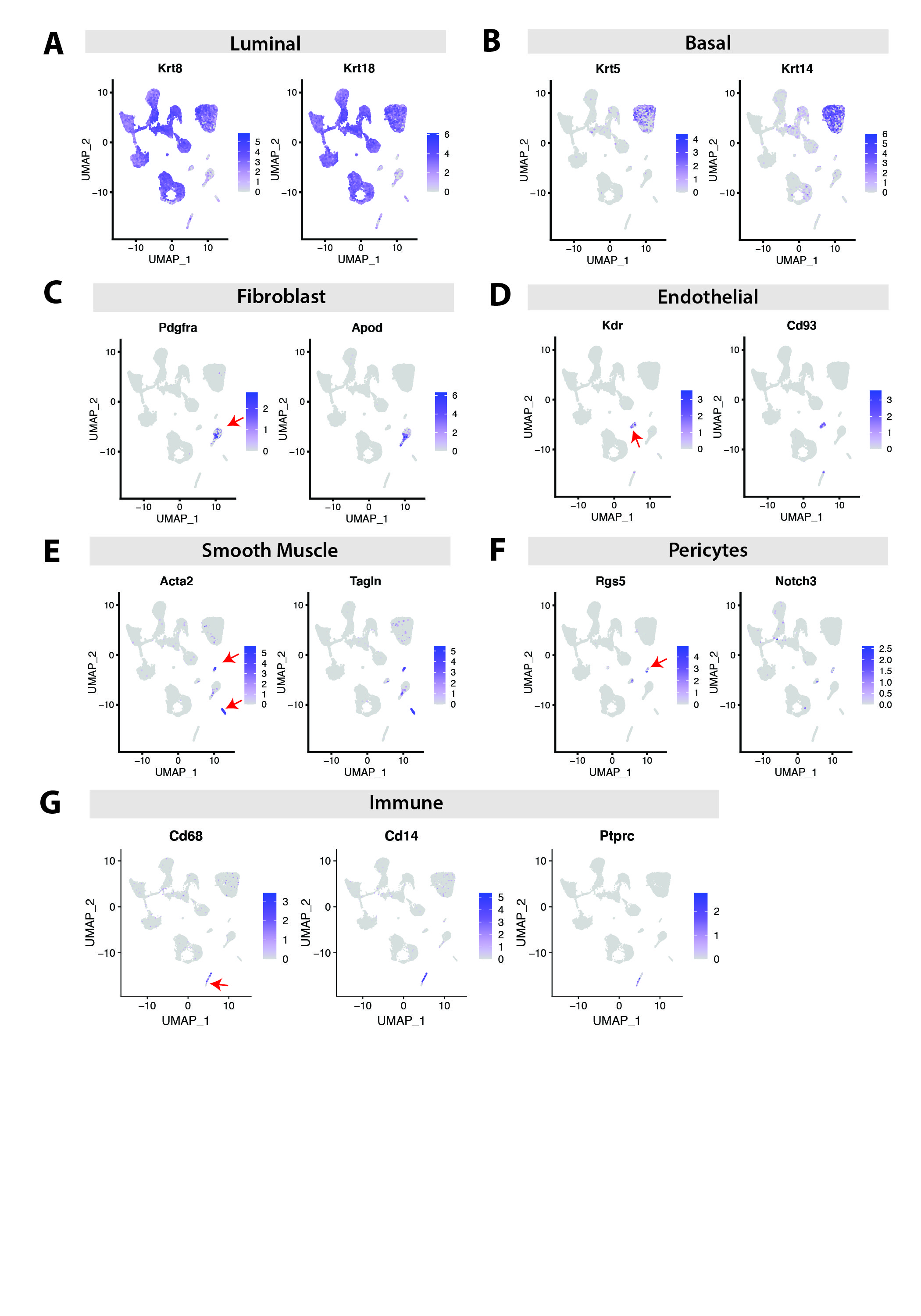

### Figure S2

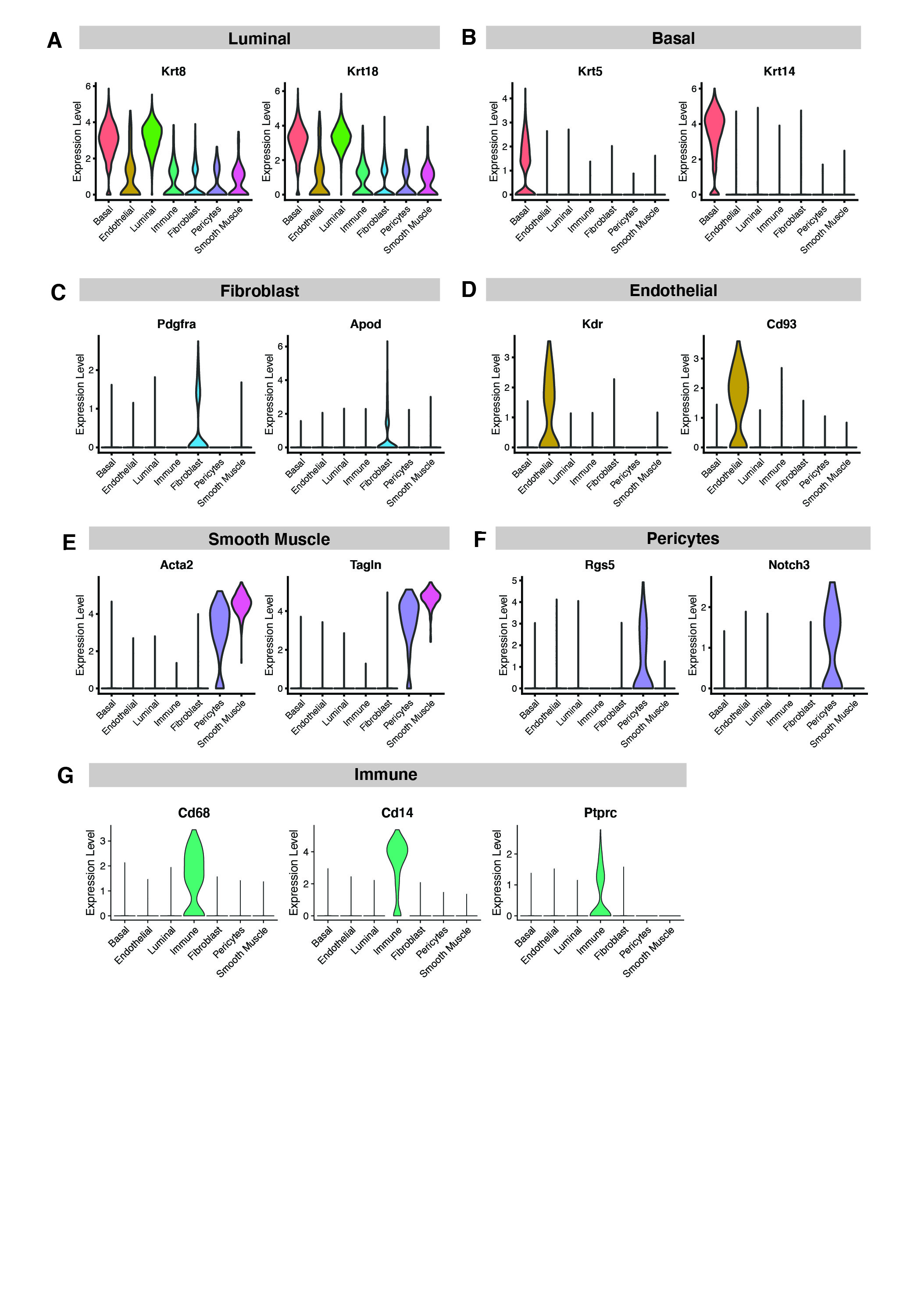

### Figure S3

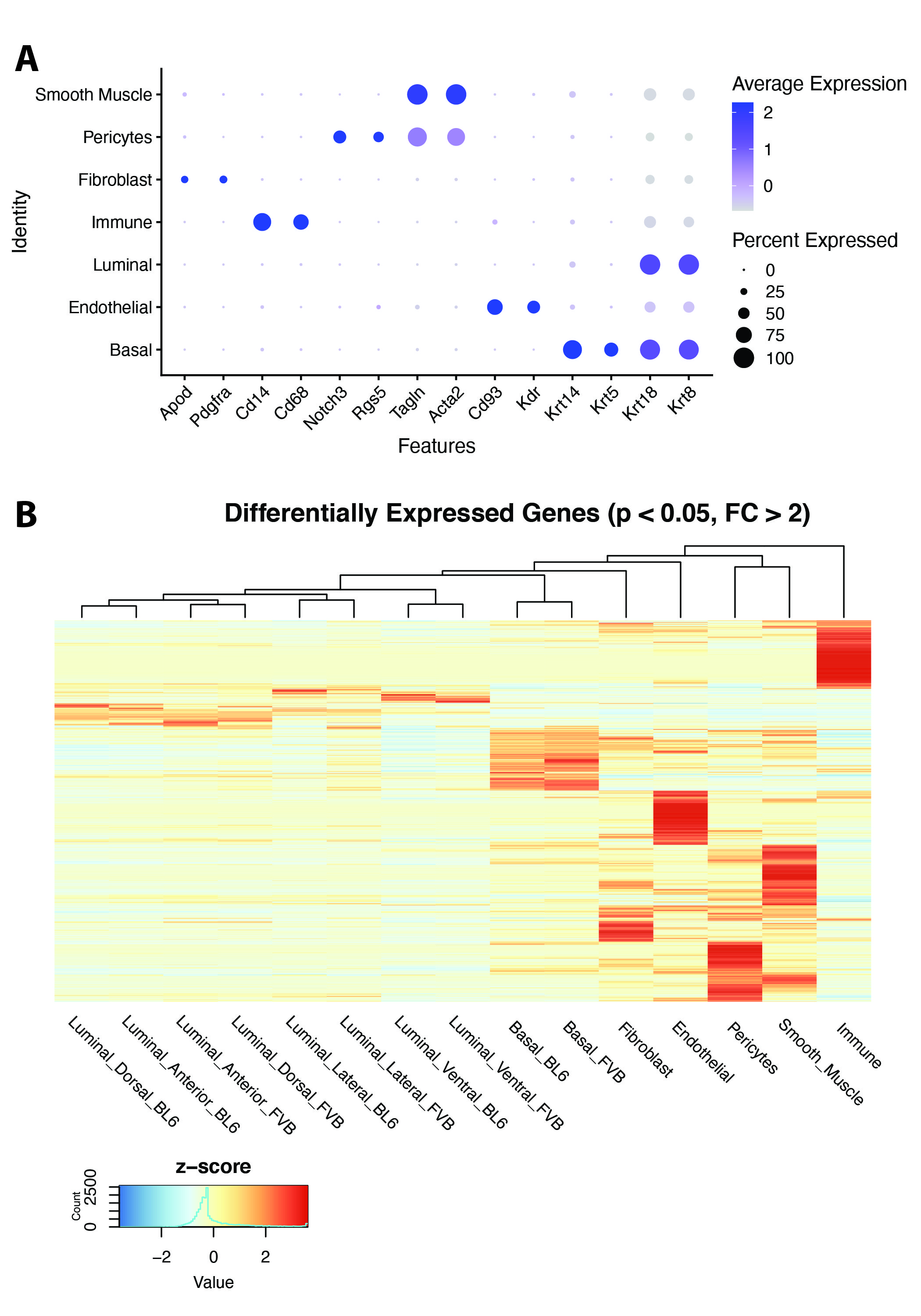

### Figure S4

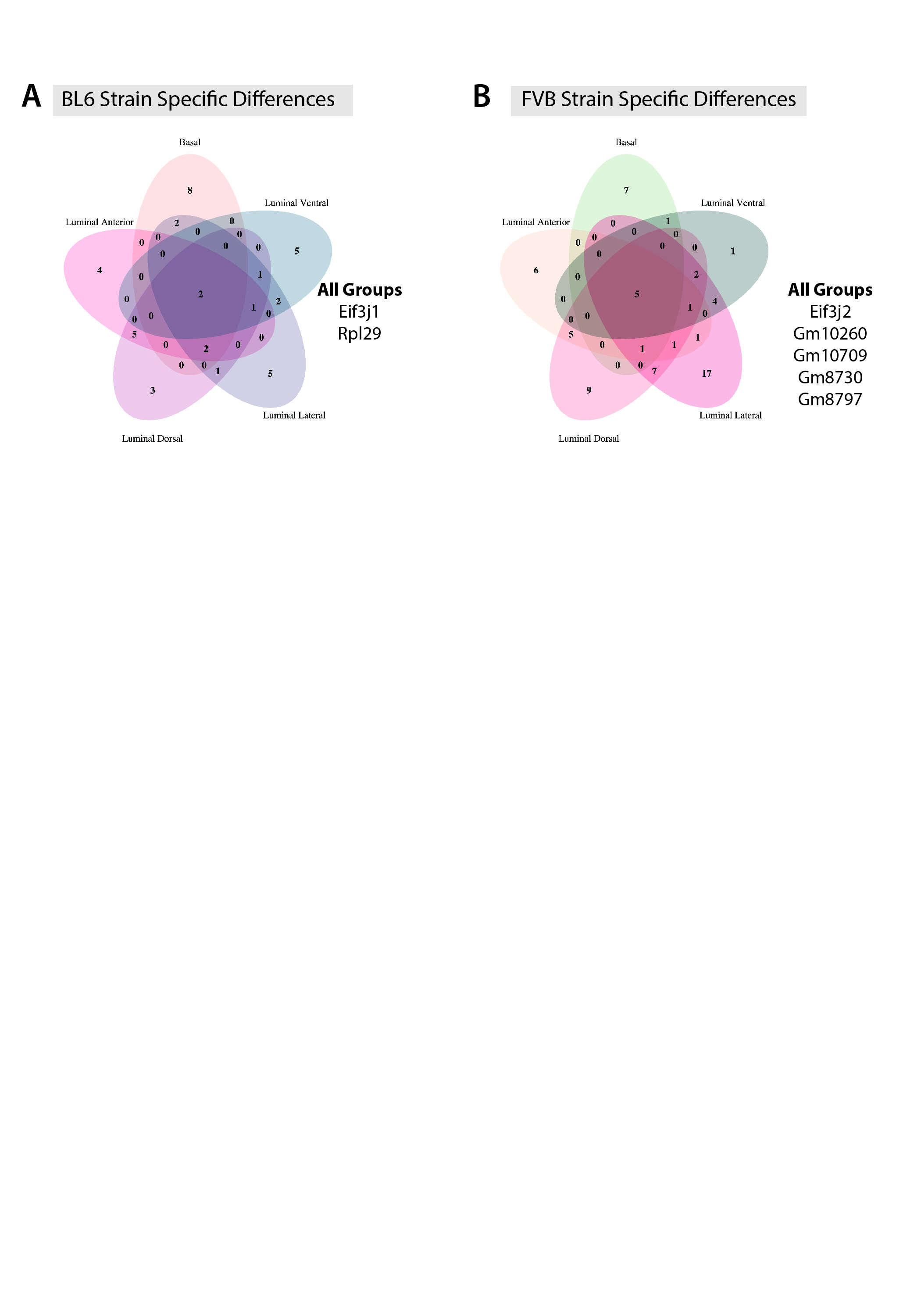

### Figure S5

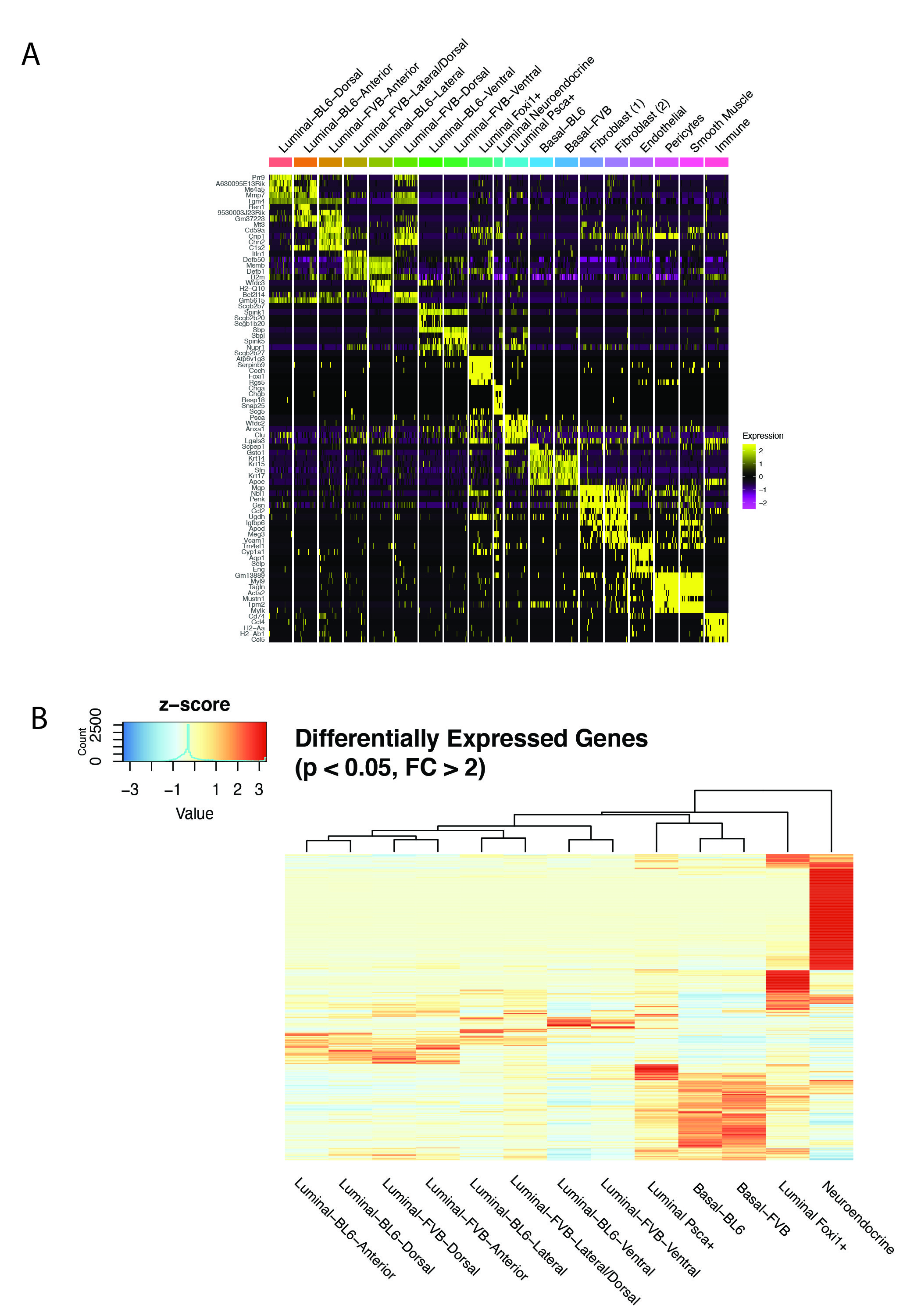

### Figure S7

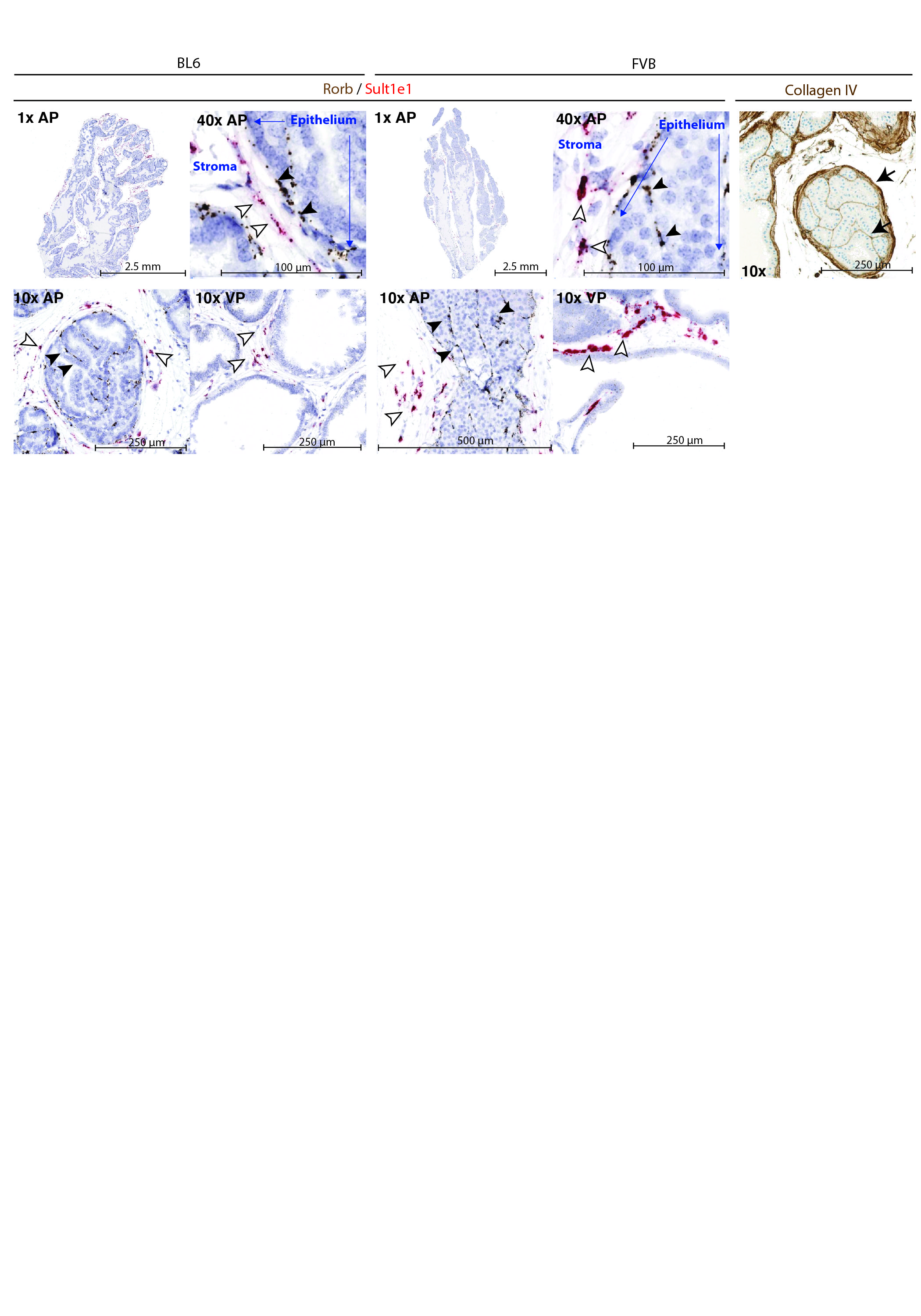

### Sigure S6

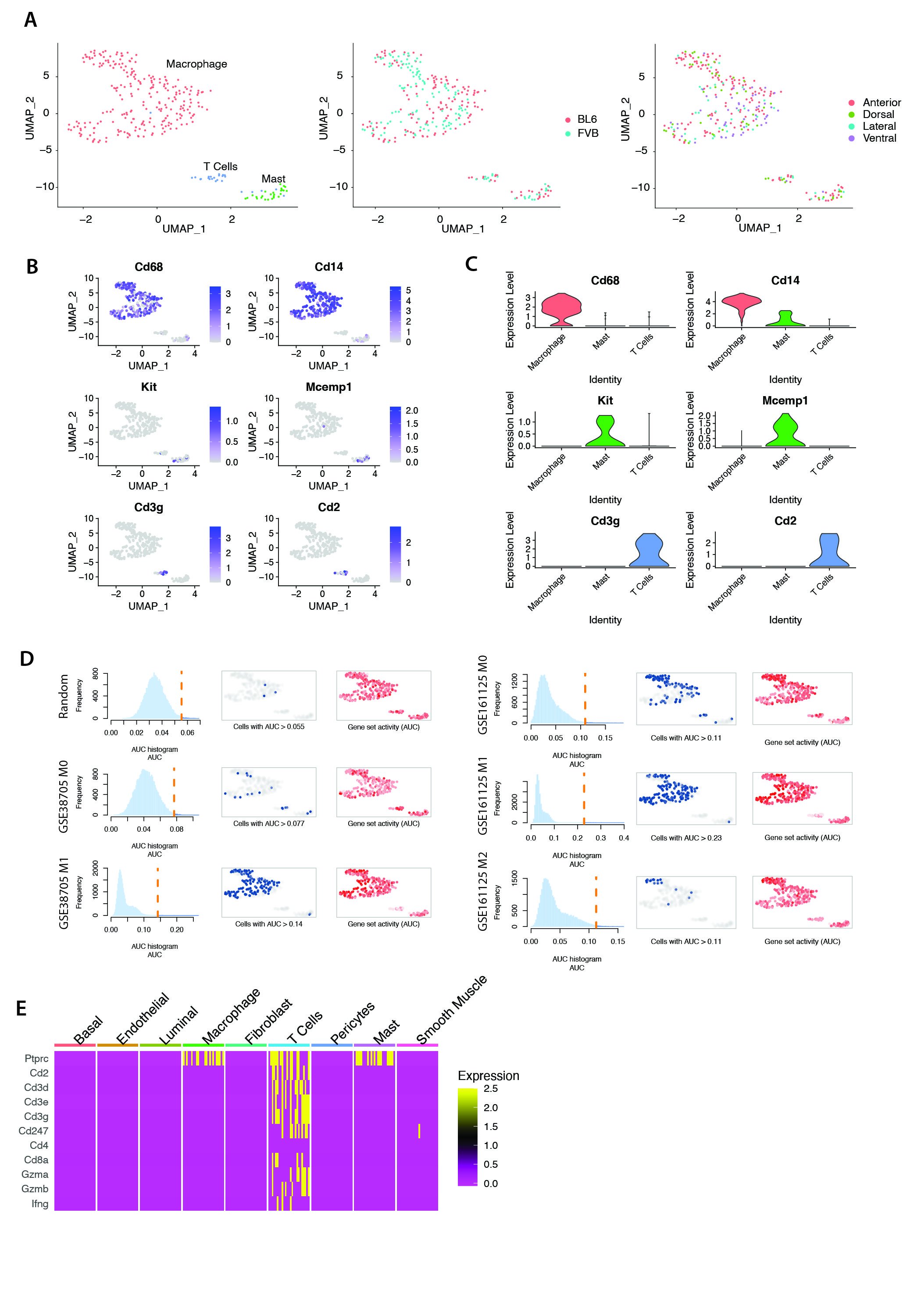
